# Site pleiotropy of a stickleback tooth and fin enhancer

**DOI:** 10.1101/2022.05.13.491895

**Authors:** Alyssa J. Rowley, Tyler A. Square, Craig T. Miller

## Abstract

Development and regeneration are orchestrated by gene regulatory networks that operate in part through transcriptional enhancers. Although many enhancers are pleiotropic and are active in multiple tissues, little is known about whether enhancer pleiotropy is due to 1) site pleiotropy, in which individual transcription factor binding sites (TFBS) are required for activity in multiple tissues, or 2) multiple distinct sites that regulate expression in different tissues. Here, we investigated the pleiotropy of an intronic enhancer of the stickleback *Bone morphogenetic protein 6* (*Bmp6*) gene. This enhancer was previously shown to regulate evolved changes in tooth number and tooth regeneration, and is highly pleiotropic, with robust activity in both fins and teeth throughout embryonic, larval, and adult life. We tested the hypothesis that the pleiotropy of this enhancer is due to site pleiotropy of an evolutionarily conserved predicted Foxc1 TFBS. Transgenic analysis and site-directed mutagenesis experiments both deleting and scrambling this predicted Foxc1 TFBS revealed that the binding site is required for enhancer activity in both teeth and fins throughout embryonic, larval, and adult development. Collectively these data support a model where the pleiotropy of this *Bmp6* enhancer is due to site pleiotropy and this putative binding site is required for enhancer activity in multiple anatomical sites from the embryo to the adult.

## Introduction

Eukaryotic gene expression is governed by enhancers, non-coding cis-regulatory elements that positively regulate transcription. Enhancers bind transcription factors that promote cell-type-specific gene expression programs throughout development and drive transcription in response to stimuli (Levine et al., 2014). Decades of genetic studies have revealed that most human disease phenotypes (Farh et al., 2015; Maurano et al., 2012), and much of natural phenotypic variation in animals are associated with sequence changes in non-coding DNA (Rebeiz & Tsiantis, 2017). These findings suggest that enhancers play a critical role in both evolution and disease, thereby motivating further study of basic enhancer biology.

Many enhancers are known to function in multiple distinct tissues, suggesting that these elements are pleiotropic. A study on chromatin from a subset of human tissues found that at least 1% of identified *cis*-regulatory elements were active in at least two spatially distinct domains (Singh & Yi, 2021). In vivo studies in fish and mice have additionally identified enhancer sequences that drive gene expression in more than one tissue type (Cleves et al., 2018; Erickson et al., 2015; Jackman & Stock, 2006; Jumlongras et al., 2012; Stepaniak et al., 2021). Two different described mechanisms of enhancer pleiotropy are (1) tightly clustered, but different, transcription factor binding sites (TFBS) that drive expression in different tissues and (2) site pleiotropy, where a single TFBS drives expression in multiple tissues (Preger-Ben Noon et al., 2018). These two mechanisms of enhancer pleiotropy have different implications for evolution; site pleiotropy would constrain evolution to preserve the integrity of critical regulatory sequences, as a single mutation in the TFBS could have a significant impact on gene expression in multiple tissues (Boffelli et al., 2004; Fish et al., 2017; Infante et al., 2015; Sabarís et al., 2019). In contrast, tightly clustered TFBS could allow for more modular evolution of gene expression in different tissues or at different developmental stages.

Threespine stickleback fish (*Gasterosteus aculeatus*) are a powerful system to study enhancer biology, as abundant natural variation throughout their adaptive radiation occurs largely due to non-coding variation, pointing to the importance of enhancers in phenotypic evolution (Jones et al., 2012). For example, changes in tooth number have evolved repeatedly in sticklebacks, with derived freshwater populations having increases in tooth number and tooth regeneration rates relative to ancestral marine populations (Cleves et al., 2014; Ellis et al., 2015). Quantitative trait loci mapping revealed a large effect locus that contains the gene *Bone Morphogenetic Protein 6* (*Bmp6*), which is dynamically expressed in both epithelial and mesenchymal cells within developing teeth (Cleves et al., 2014). Further high-resolution genetic mapping of this large effect locus linked the evolved phenotypic effects to an enhancer in the fourth intron of *Bmp6* (Cleves et al., 2018). This intronic enhancer is highly pleiotropic and drives expression in all developing and regenerating teeth, as well as the distal edges of the pectoral, median, and caudal fins (Cleves et al., 2018). Comparing the marine and freshwater versions of this intronic enhancer in doubly transgenic lines revealed evolved spatial shifts in enhancer activity in both tooth epithelium and mesenchyme, consistent with an evolved change in enhancer activity driving phenotypic evolution of tooth number (Stepaniak et al., 2021). In addition, although both enhancers drive strong expression in the distal edges of embryonic fins, only the freshwater enhancer was detected in fin ray joints in pectoral and caudal fins (Stepaniak et al., 2021). However, whether the enhancer utilizes common inputs for tooth and fin activity, and which sites are required for this activity are unknown.

Previous studies have found that fish teeth and mammalian hair share many aspects of early development: both are derived from placodes (Pispa & Thesleff, 2003), continuously regenerate in adults, are regulated by BMP signaling during replacement (Jia et al., 2013; Vainio et al., 1993; Wang et al., 2012), and express similar batteries of genes during regeneration (Square et al., 2021). In mice, BMP signaling has been proposed as a major regulator of maintaining stem cell quiescence. Conditional ablation of the *Bmpr1a* gene in skin epithelium activated the stem cell niche, causing quiescent hair follicle stem cells to proliferate (Kandyba et al., 2013; Kobielak et al., 2007). Further, Foxc1 regulates *Bmp6* expression in regenerating hair. Foxc1 binds to a regulatory region adjacent to *Bmp6* in mice and inhibits hair regeneration, as conditional knockout of *Foxc1* in skin results in accelerated hair regeneration (Wang et al., 2016). Moreover, Foxc1 was proposed to be a candidate regulator of the *Bmp6* intron 4 enhancer, as a putative Foxc1 TFBS was identified within this enhancer (Cleves et al., 2018). Given that this enhancer is expressed in regenerating teeth, has been linked to evolved changes in tooth regeneration, and the similarities between hair and tooth regeneration, a parsimonious model is that BMP signaling negatively regulates mammalian hair and fish tooth regeneration using homologous gene regulatory networks. Here we test the hypotheses that (1) a predicted Foxc1 TFBS is required for *Bmp6* enhancer activity in developing and regenerating teeth and (2) this predicted Foxc1 TFBS has site pleiotropy and is required for enhancer activity in teeth and fins.

## Methods

### Animal husbandry

All animal work was approved by UC Berkeley IACUC protocol AUP-2015–01-7117. Sticklebacks were raised as previously described (Cleves et al., 2014; Square et al., 2021). All experiments used lab-reared marine (Rabbit Slough, Alaska) sticklebacks.

### Generation of transgenic GFP enhancer stickleback lines

*Tol2* plasmid transgenesis was performed using pT2HE plasmid backbone as previously described (Erickson et al., 2016; O’brown et al., 2015). To generate GFP reporter constructs, PCR-based site-directed mutagenesis was done on plasmids containing the ∼1300 bp Paxton benthic (PAXB) high-tooth-associated intron four enhancer allele (Cleves et al., 2018; Stepaniak et al., 2021). Primers for site-directed mutagenesis were designed using the Quickchange tool and PCR was carried out according to Erickson et al. (Erickson et al., 2015) https://www.agilent.com/store/primerDesignProgram.jsp. Primer sequences used for the deleted were CTAATTACCCCGACGAGGTCGGGTGGGGAG and CTCCCCACCCGACCTCGTCGGGGTAATTAG and for the scrambled CCCCACCCGACCTTTTGAATACCGTCGGGGTAATTA and TAATTACCCCGACGGTATTCAAAAGGTCGGGTGGGG. Transposase messenger RNA was synthesized as described previously (Kawakami & Shima, 1999). Stickleback embryos at the one-cell stage were microinjected as described (Erickson et al., 2016). Two stable transgenic lines were generated with each of the intact, scrambled, and deleted reporter constructs.

### Imaging fluorescent transgenic sticklebacks

Transgenic GFP reporter lines were imaged using a Leica M165FC stereoscope with a DFC340 FX camera. Transgenic embryos were imaged live. Transgenic juvenile and adult fish were fixed in 4% paraformaldehyde for 4 hours for 9-19 mm standard length (SL) or overnight (>20mm SL). Teeth were imaged using Montage z-stack projections on a Leica M165FC dissecting microscope with a GFP2 filter.

### Bioinformatics

The following genome assemblies were used to compare *Bmp6* sequences: stickleback *Gasterosteus aculeatus*, “Gac”: Broad/gasAcu, medaka *Oryzias latipes*, “Ola”: NIG/UT MEDAKA1/oryLat2, zebrafish *Danio rerio*, “Dre”: GRCz10/danRer10, gar *Lepisosteus oculatus* “Loc”: LepOcu1 (GCA_000242695.1). A ∼25 kb window centered on stickleback *Bmp6* was aligned to orthologous sequences from these three other species using mVISTA and LAGAN at https://genome.lbl.gov/vista/index.shtml (Frazer et al., 2004) using default visualization parameters of 100 bp windows showing minimum y-axis of 50% and 70% sequence identity to color windows as conserved. The entire fourth intron sequence from all four species was aligned as above, and the first 5.5 kb of the ∼6.5 kb intron shown in Figure 2C. Sequences orthologous to the minimally sufficient ∼500 bp *Bmp6* intron 4 enhancer (Cleves et al., 2018) were aligned using Clustal Omega at https://www.ebi.ac.uk/Tools/msa/clustalo/ (Sievers et al., 2011) and default parameters. The resulting text alignment was opened in MS Word and conserved bases in all four species highlighted manually.

### In situ hybridizations on sections

Sample preparation, sectioning, *in situ* hybridization, and riboprobe synthesis were carried out as previously described (Square et al., 2021). Riboprobes were designed as described previously (Ellis et al., 2016) against *Foxc1a* and *Foxc1b*. Riboprobe template plasmids were created by PCR cloning gene fragments from genomic DNA using primers to *Foxc1a*: 5’-GCCGctcgagGGGACAGGTCTAGCCACTTG-3’; 5’-GCCGtctagaACGGCGATA-TACACGTTCCT-’3 and *Foxc1b*: 5’-GCCGctcgagCCTGCCCGACTATTGCATCA-3’; 5’-GCCGactagtAGACGGCAC-TTTATTAAACAAACA-3’.

## Results

Previous research mapped evolved increases in tooth number in freshwater sticklebacks to an enhancer located in the fourth intron of *Bmp6* (Cleves et al., 2014, 2018). This enhancer drives robust expression in all developing pharyngeal and oral teeth, as well as in the distal margins of the pectoral and median fins (Fig. 1A,B) (Cleves et al., 2018; Stepaniak et al., 2021). The sequence of this intronic enhancer is evolutionarily conserved in teleosts, with clear homology detectable in medaka and zebrafish, as well as to gar, an outgroup to teleosts (Fig. 1C,D). Comparison of enhancer sequences within fish and outgroups suggests different levels of constraint within this enhancer, possibly representing functionally required TFBSs. Of those regions with 100% sequence conservation across the aforementioned fish species, one includes a nine base pair motif TTTGTTTAC that perfectly matches the binding site consensus previously reported for human FOXC1 (Fig. 1D, Fig. S1) (Berry et al., 2008).

**Figure 1.**
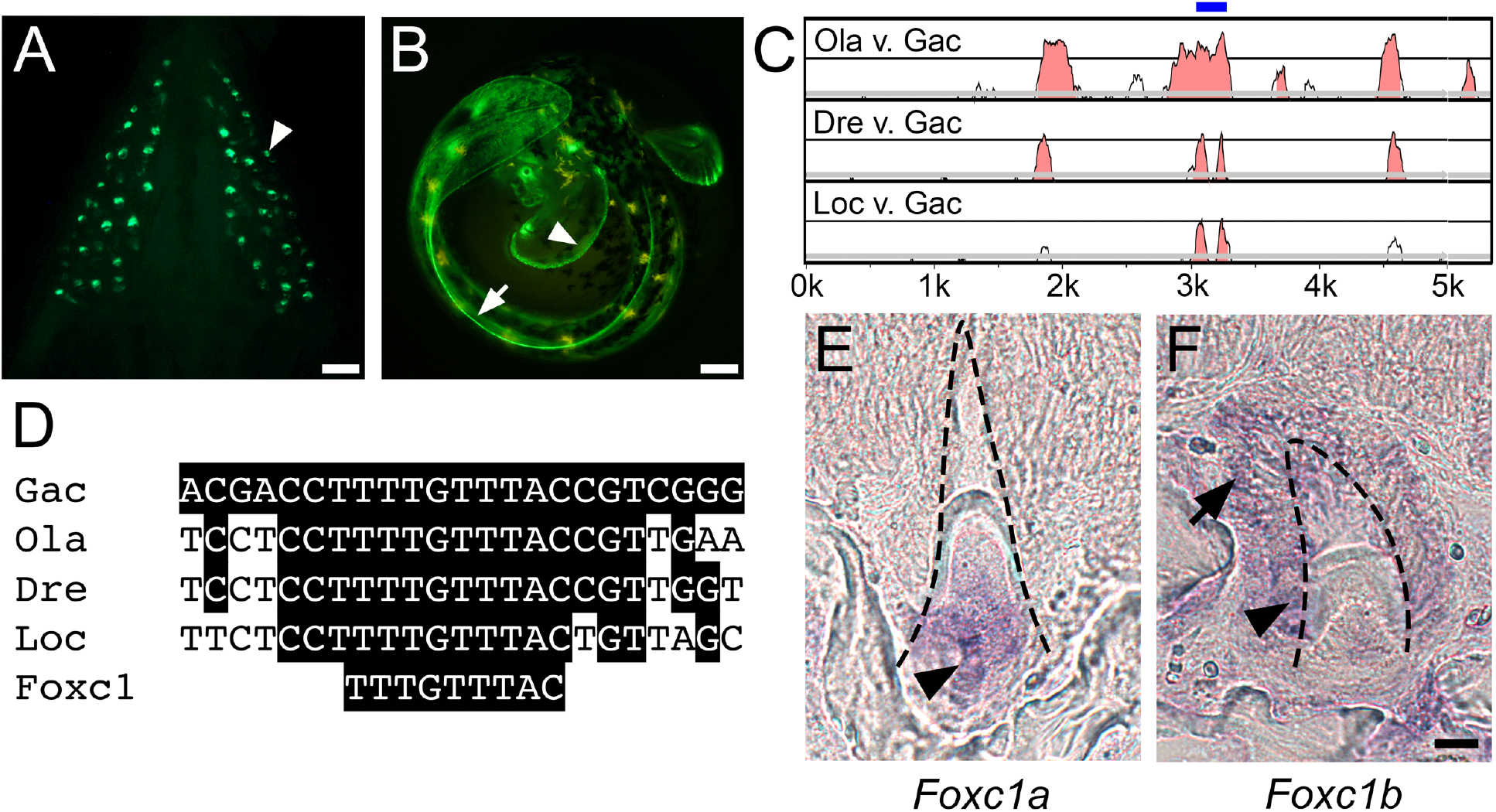
A pleiotropic tooth and fin enhancer includes an evolutionarily conserved predicted Foxc1 binding site and Foxc1 genes are expressed in teeth. eGFP reporter gene driven by a 1.3 kb *Bmp6* intronic enhancer (Square et al., 2021; Stepaniak et al., 2021) reveals expression in (A) ventral pharyngeal teeth (arrowhead) of juvenile stickleback fish and (B) distal edge of pectoral (arrowhead) and median (arrow) fin fold of unhatched 7dpf developing stickleback embryo. (C) mVISTA LAGAN sequence alignment showing central region of stickleback Bmp6 intron four is conserved to gar. Intron four of stickleback *Bmp6* (x-axis) is aligned to medaka (top), zebrafish (middle) and gar (bottom). Graph shows a sliding window of 100 base pairs, with y-axis starting at 50% identity and windows shaded at a threshold of 70% nucleotide identity. Blue bar above highlights central island of conservation (see Figure S1). Gac = *Gasterosteus aculeatus* (stickleback), Ola = *Oryzias latipes* (medaka), Dre = *Danio rerio* (zebrafish), Loc = *Lepisosteus oculatus* (gar). (D) Predicted Foxc1 binding site TTTGTTTAC (Berry et al., 2008) within central island of conservation is conserved from sticklebacks to gar. In situ hybridization of *Foxc1a* (E) and *Foxc1b* (F) expression on sections of adult stickleback pharyngeal teeth. *Foxc1a* expression was detected in dental mesenchyme (black arrowhead), while *Foxc1b* expression was detected in the inner (black arrowhead) and outer dental epithelium (black arrow). Black dotted lines demarcate the outline of the developing mineralized tooth. Scale bar = 10 µm.

Teleosts underwent a whole genome duplication (Amores et al., 1998), resulting in pairs of co-orthologs of many teleost genes relative to outgroups. In sticklebacks, both *Foxc1a* and *Foxc1b* have been maintained. Expression of both *Foxc1* genes was detected in developing adult teeth, with *Foxc1a* largely restricted to the dental mesenchyme (Fig. 1E), and *Foxc1b* distributed throughout the inner and outer dental epithelium (Fig. 1F). Thus, *Foxc1* gene expression overlaps with activity of the *Bmp6* intronic enhancer in teeth (Stepaniak et al., 2021).

To test the hypothesis that the predicted Foxc1 TFBS is required for enhancer activity, we introduced mutations in a *Bmp6* reporter construct previously described (Cleves et al., 2018; Stepaniak et al., 2021). The unaltered construct drove GFP expression in several tissues including the pectoral and median fins, and oral and pharyngeal teeth. Site directed mutagenesis was used to make two mutant enhancer variants: (1) a scrambled enhancer, where two thymidine nucleotides in the predicted Foxc1 TFBS were mutated to adenine, and (2) a deleted enhancer, where the entire nine base pair Foxc1 predicted TFBS was deleted (Fig. 2A). We generated two stable transgenic lines each for three different enhancer transgenes: intact wild-type, scrambled, and deleted (Fig. 2A). For all three genotypes, we imaged transgenic larvae, juveniles, and adults to test for possible effects on enhancer activity in different tissues. All scrambled and deleted transgenes dramatically reduced GFP expression in the pectoral and median fins of the developing embryo (Fig. 2B).

**Figure 2.**
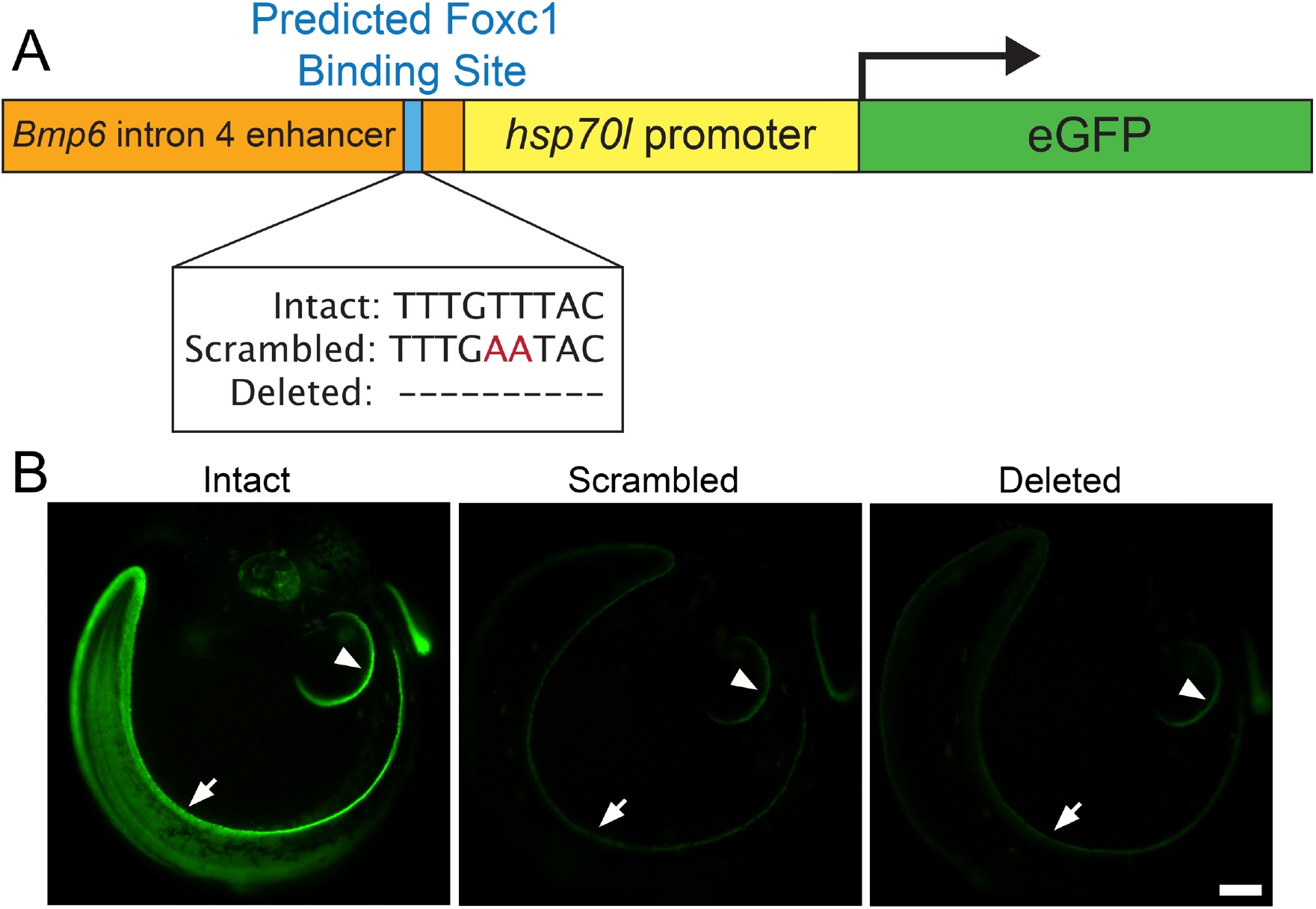
Predicted Foxc1 binding site is required for embryonic enhancer activity. (A) Schematic of eGFP reporter construct with *Bmp6* intronic enhancer with intact, scrambled, and deleted sequences for the predicted Foxc1 binding site upstream of the *hsp70l* promoter driving eGFP. (B) eGFP reporter expression pattern driven by the *Bmp6* intronic enhancer with intact, scrambled, or deleted sequence for the Foxc1 binding site in 7 dpf (stage 22, (Swarup, 1958)) stickleback embryos. Strong expression was seen in the distal edge of the developing pectoral fin (arrowhead) and median fin (arrow) for the enhancer with an intact sequence of the predicted Foxc1 binding site (left). Severely reduced eGFP expression in all domains of enhancer activity was detected for enhancers with the scrambled (middle) or deleted (right) sequences of the predicted Foxc1 binding site. Scale bar = 100 µm.

While the stable transgenic reporter lines containing the intact binding site drove robust expression in every detectable tooth in the pharyngeal and oral jaw, neither the scrambled nor deleted reporter lines drove detectable GFP expression in pharyngeal teeth (Fig. 3). As a control, GFP lens expression driven by the *hsp70l* promoter (Erickson et al., 2015) was comparably bright in the intact, scrambled, and deleted transgenic lines (Fig. 3A). Within developing primary oral and pharyngeal teeth, transgene GFP expression was observed in the dental epithelium and mesenchyme for the intact enhancer, but in neither epithelium nor mesenchyme for the scrambled or deleted enhancer in pharyngeal teeth (Fig. 3). GFP expression was also seen in the epithelium and mesenchyme of adult teeth for the intact enhancer. Faint GFP expression was detected in some oral teeth in the premaxilla at adult stages for the scrambled enhancer (Fig 3C). Overall, adult tooth expression was largely abolished in the scrambled and deleted enhancer reporter lines, suggesting that the predicted Foxc1 TFBS is required for enhancer activity in replacement as well as primary teeth (Fig. 3).

**Figure 3.**
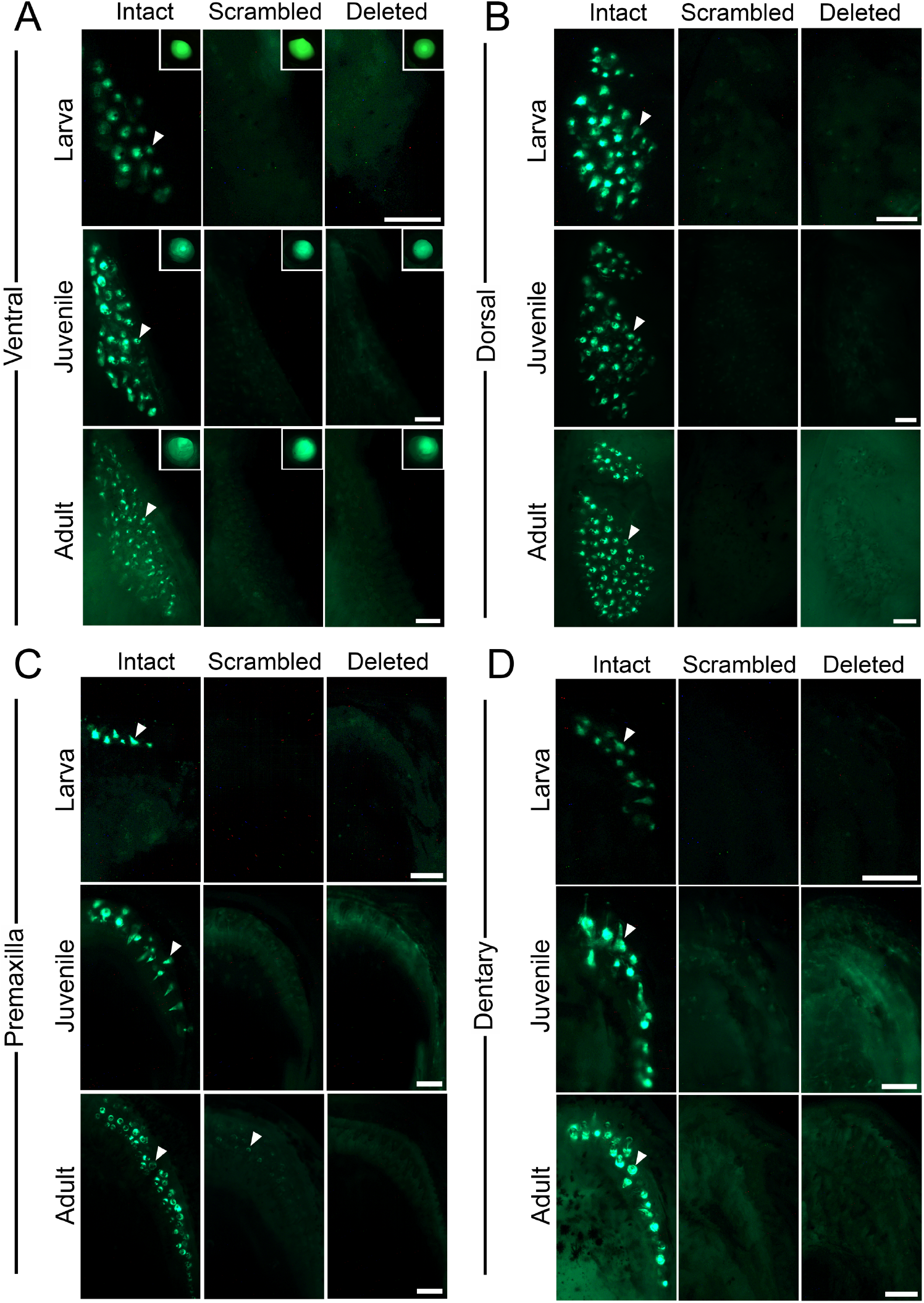
Predicted Foxc1 binding site is required for pharyngeal and oral tooth expression in larva, juveniles, and adults. eGFP reporter gene expression in ventral (A,C) and dorsal (B,D) pharyngeal (A,B) and oral (C,D) jaws in larva (7-11mm SL), juvenile (17-21mm SL), and adult (37-55mm SL) sticklebacks. GFP expression was detected in the epithelium and mesenchyme of developing teeth (arrowheads) in larva, juvenile and adult stages with the intact sequence of the predicted Foxc1 binding site (left columns). No expression was detected in any developing teeth of larva, juvenile, and adults with the scrambled or deleted sequence of the predicted Foxc1 binding site (middle and right columns, respectively), except for faint expression in some adult premaxilla teeth (arrow in adult premaxilla panel in C). (A insets) Lens of eye with eGFP expression as an internal control driven by the *hsp70l* promoter used in the transgenic construct. Scale bars = 100 µm (larva and juvenile), 250 µm (adult).

Additionally, we found that the predicted Foxc1 TFBS is required for enhancer function in fins at all developmental time points examined, including in the intersegmental joints of the juvenile and adult pectoral and caudal fins. All fin domains were nearly abolished in both the scrambled and deleted reporter lines at all developmental stages (Fig. 4). GFP expression was detected in rare intersegmental joints of the pectoral fins in juveniles for the scrambled enhancer, and in the base of the pectoral fin rays for both the scrambled and deleted construct (Fig. 4A). Some GFP expression persisted in the caudal peduncle for the deleted enhancer (Fig. 4B). In general, all sites of fin expression, like tooth expression, were severely reduced in both the scrambled and deleted enhancer lines relative to the intact enhancer lines. These data support a crucial role for this single putative Foxc1 binding site in positively regulating both tooth and fin expression across development from embryos to adults.

**Figure 4.**
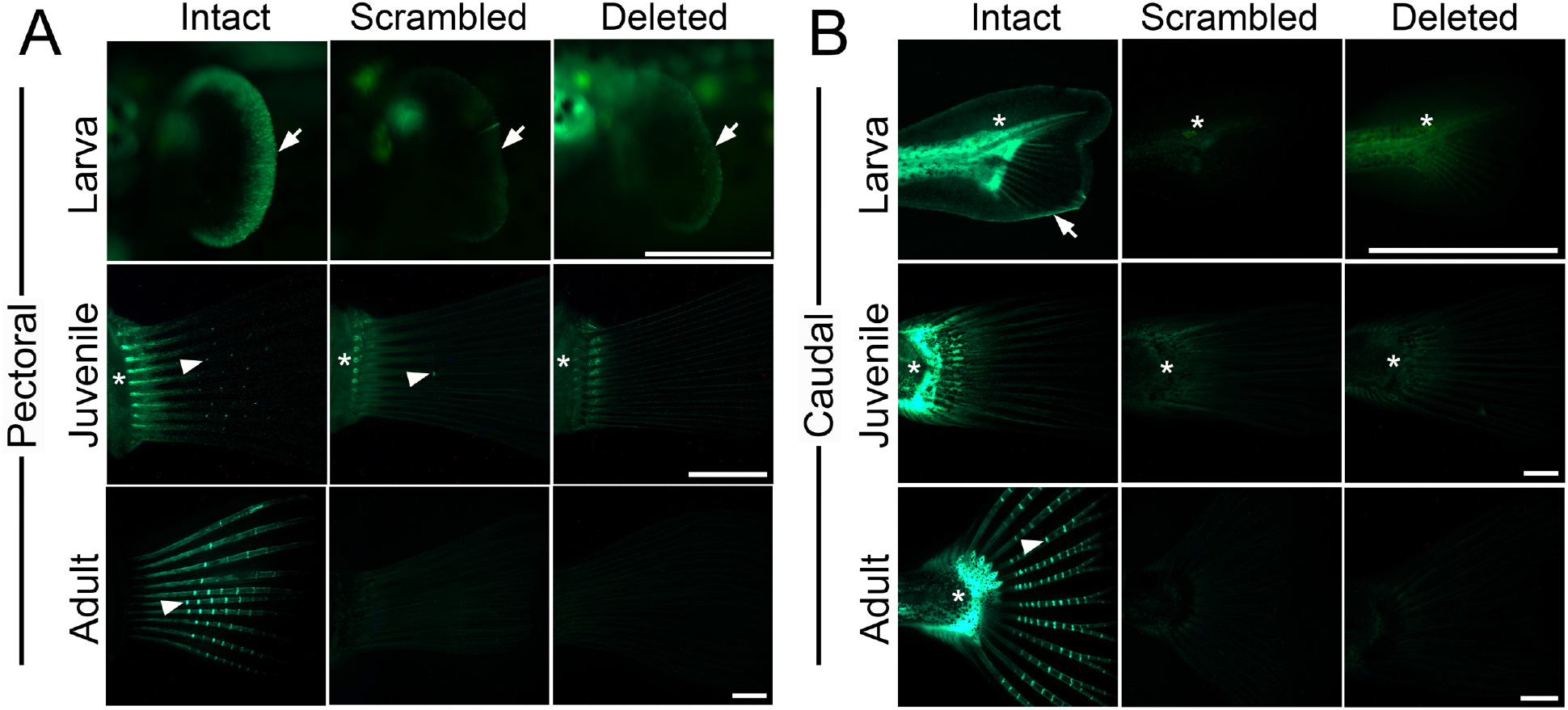
Predicted Foxc1 binding site is required for fin expression in larva, juveniles, and adults. eGFP reporter gene expression in the left pectoral fin (A) and caudal fin (B) of larva, juveniles and adults. (A) Expression was detected in the distal edges of the pectoral fin (arrows) in larva, the base of the pectoral fin rays (asterisk), and the intersegmental joints (arrowheads) of juveniles and adults with the intact sequence of the predicted Foxc1 binding site (left column). Severely reduced expression was seen in the distal edges of the pectoral fin in larva and the intersegmental joints of juveniles and adults with the scrambled or deleted sequence of the predicted Foxc1 binding site (middle and right columns, respectively), while some expression at the base of the fin rays remained. (B) eGFP reporter expression in the caudal fin. In larva, expression was observed in the distal edge of the median fin fold (arrow) and caudal peduncle (asterisk) with the intact enhancer, but was severely reduced with the scrambled and deleted enhancers. Some peduncle expression remained with the deleted enhancer (right). In juveniles and adults, expression was detected in the peduncle (asterisk) and the intersegmental joints of the caudal fin (arrowheads) for the intact enhancer, but not for the scrambled or deleted enhancer. Scale bars = 500 µm (larva), 1 mm (juvenile), 2 mm (adult).

## Discussion

### A short, conserved, pleiotropic binding site is required for enhancer activity in teeth and fins

Here we show a predicted Foxc1 TFBS is required for *Bmp6* intron four enhancer activity in teeth and fins at all stages examined of stickleback development. Altering the predicted Foxc1 TFBS by scrambling two nucleotides or deleting nine nucleotides via site-directed mutagenesis severely downregulated enhancer activity in both teeth and fins, suggesting that this predicted TFBS is critical for enhancer function in multiple expression domains. This site pleiotropy is reminiscent of a previously described enhancer 5’ to the stickleback *Bmp6* gene, where a conserved predicted SMAD3 binding site is required for enhancer activity in teeth and fins (Erickson et al., 2015). While both marine and freshwater intron four *Bmp6* enhancers drive strong expression in the distal edges of embryonic fins, only the freshwater enhancer was detected in intersegmental joints in pectoral and caudal fins (Stepaniak et al., 2021). The intact enhancer tested here was the freshwater allele and drove robust fin ray joints as previously reported. Both early distal fin fold and later fin joint expression domains driven by the intact enhancer were largely absent in the scrambled and deleted Foxc1 TFBS reporter lines.

Enhancers have displayed both site pleiotropy and tightly linked binding sites driving expression in different tissues at different developmental stages. A study investigating the *shavenbaby* enhancers in *Drosophila* found that while one enhancer used the same TFBS to regulate embryonic and pupal expression (site pleiotropy), another enhancer required distinct tightly linked binding sites for expression in different tissues (Preger-Ben Noon et al., 2018). While we hypothesize that this predicted TFBS studied here is bound by Foxc1, it may serve as a binding site for other Fox or non-Fox transcription factors with similar binding site affinity. It is possible that in generating scrambled or deleted TFBSs, one or more new TFBS were generated. However, it would be unlikely that both deleted and scrambled nucleotides would create the same TFBS, given the different primary sequence of these two mutations. Once stickleback Foxc1 antibodies are available, biochemical experiments could test whether stickleback Foxc1a and/or Foxc1b binds this predicted TFBS. Genetic experiments, crossing this enhancer into *Foxc1a* and *Foxc1b* single and double mutants could directly test whether either or both of these duplicate *Foxc1* genes regulate this pleiotropic enhancer.

### Roles for Foxc1 in BMP regulation

Previous studies have revealed *Foxc1* as a molecular switch for regulating regeneration in mouse hair follicles. As a transcription factor, Foxc1 activates BMP signaling in self-renewing hair follicle stem cells (HFSCs) (Wang et al., 2016). The conditional knockout of *Foxc1* in mouse results in reduced quiescence in HFSCs and an increase in hair regeneration rates. RNA sequencing (RNA-Seq) and quantitative polymerase chain reaction (qPCR) revealed that genes associated with quiescence of HFSCs, like *Bmp6*, were down-regulated. The direct regulation of *Bmp6* by Foxc1 was supported by chromatin immunoprecipitation PCR (Wang et al., 2016). Here we suggest Foxc1 regulates the BMP pathway for tooth, and possibly fin, development and regeneration through this TFBS in *Bmp6*. Future experiments analyzing the phenotypic effects of *Foxc1a* and *Foxc1b* single and double mutants could shed light on the role that these stickleback *Foxc1* co-orthologs play in maintaining quiescence and organ development and regeneration through the BMP pathway.

### Foxc1 in organogenesis and chromatin remodeling

Previous studies of enhancer evolution have suggested that enhancers are born as “proto-enhancers” with an initial low-information content but containing a TFBS that serves as a nucleation point around which other TFBSs then evolve (Emera et al., 2016). The winged-helix structure of the forkhead DNA-binding domain, which is highly conserved across all members of the Fox family, resembles the structure of the linker histone H1, a key modulator of chromatin structure. Because of its structure and chromatin binding interactions, FoxA proteins have been proposed to act as pioneer transcription factors, opening compacted chromatin to allow the binding of other transcription factors (Cirillo et al., 2002; Lee et al., 2005). Foxd3 has been shown to act as a pioneer factor by re-arranging the chromatin landscape and opening *cis*-regulatory elements to maintain multipotency in the early neural crest lineage (Lukoseviciute et al., 2018). Together, these studies suggest Fox family transcription factors promote chromatin accessibility within promoter and enhancer regions.

The loss of Foxc1 co-orthologs (*foxc1a* and *foxc1b*) in zebrafish results in severe reductions of upper facial cartilages as well as missing trabecular cartilage of the neurocranium. The zebrafish *foxc1a*^*−/−*^ mutants died by 7 dpf, and displayed facial cartilage defects and *foxc1b*^*−/−*^ mutants displayed a truncation in the symplectic cartilage (Xu et al., 2018). In double homozygous *foxc1a*; *foxc1b* mutants, severe reductions of upper facial cartilages occurred, suggesting that Foxc1a and Foxc1b act redundantly in upper facial cartilage development (Xu et al., 2018). In zebrafish embryos doubly homozygous *foxc1a* and *foxc1b* mutations, the cartilaginous skeleton did not form properly, likely because many of the regulatory sequences in cartilage regulatory genes were not accessible (Xu et al., 2021). These results support a model in which zebrafish Foxc1 proteins serve as pioneer transcription factors to open chromatin, making it accessible for other transcription factors to bind and regulate genomic regions responsible for cartilage development. It is possible that stickleback Foxc1 proteins may serve similar roles by opening chromatin regions specific to tooth and fin enhancers so other transcription factors can regulate development or regeneration. Future experiments analyzing the genetic effects of *Foxc1a* and *Foxc1b* single and double null-mutants could elucidate the role of the Foxc1 co-orthologs in sticklebacks, and test whether they serve as pioneer transcription factors to promote chromatin accessibility.

## Supplemental Figure

**Figure S1.**
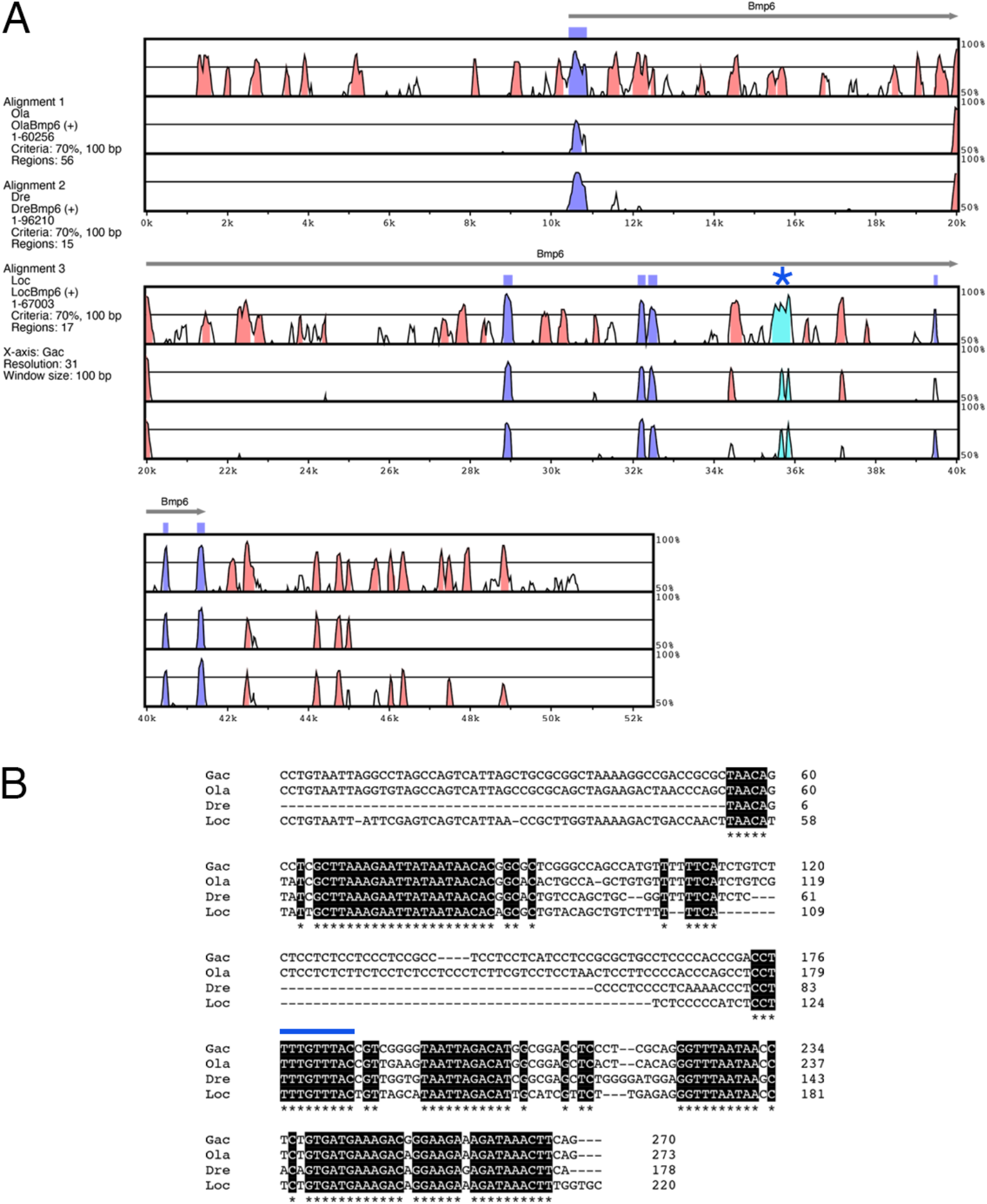
Evolutionary conservation of a predicted Foxc1 binding site within the fourth intron of Bmp6. (A) mVISTA LAGAN alignment of stickleback *Bmp6* locus to medaka, zebrafish, and gar genome sequences. Gac = *Gasterosteus aculeatus* (stickleback), Ola = *Oryzias latipes* (medaka), Dre = *Danio rerio* (zebrafish), Loc = *Lepisosteus oculatus* (gar). The x-axis shows ∼25 kb centered on stickleback *Bmp6* aligned to medaka (Ola, top), zebrafish (Dre, middle) and gar (Loc, bottom). Y-axis starts at 50% nucleotide identity and sliding windows of 100 bp are colored at a 70% nucleotide identity threshold. *Bmp6* exons are annotated as purple regions. The minimally sufficient ∼500 base pair intron 4 enhancer (Cleves et al., 2018) is evolutionarily conserved, annotated in light blue, and marked with a blue asterisk. (B) Zooming in on this conserved region reveals that a predicted Foxc1 binding site (TTTGTTTAC; Berry et al., 2008) (marked by blue line above) is perfectly conserved in medaka, zebrafish, and gar.

## Acknowledgements

We thank Mark Stepaniak and Josh Tworig for assistance and advice on developing and injecting transgenic constructs, and Sophie Archambeault, Naama Weksler, and Hernan Garcia for suggestions on the manuscript.

## Funding

This work was supported by the National Institutes of Health (DE021475 and DE027871).

## Notes

### Competing Interest Statement

The authors have declared no competing interest.

## References

Amores, A., Force, A., Yan, Y. L., Joly, L., Amemiya, C., Fritz, A., … Postlethwait, J. H. (1998). Zebrafish hox clusters and vertebrate genome evolution. Science, 282(5394), 1711–1714. https://doi.org/10.1126/science.282.5394.1711

Berry, F. B., Skarie, J. M., Mirzayans, F., Fortin, Y., Hudson, T. J., Raymond, V., … Walter, M. A. (2008). FOXC1 is required for cell viability and resistance to oxidative stress in the eye through the transcriptional regulation of FOXO1A. Human Molecular Genetics, 17(4), 490–505. https://doi.org/10.1093/hmg/ddm326

Boffelli, D., Nobrega, M. A., & Rubin, E. M. (2004, June). Comparative genomics at the vertebrate extremes. Nature Reviews Genetics. https://doi.org/10.1038/nrg1350

Cirillo, L. A., Lin, F. R., Cuesta, I., Friedman, D., Jarnik, M., & Zaret, K. S. (2002). Opening of compacted chromatin by early developmental transcription factors HNF3 (FoxA) and GATA-4. Molecular Cell, 9(2), 279–289. https://doi.org/10.1016/S1097-2765(02)00459-8

Cleves, P. A., Ellis, N. A., Jimenez, M. T., Nunez, S. M., Schluter, D., Kingsley, D. M., & Miller, C. T. (2014). Evolved tooth gain in sticklebacks is associated with a cis-regulatory allele of Bmp6. Proceedings of the National Academy of Sciences of the United States of America, 111(38), 13912–13917. https://doi.org/10.1073/pnas.1407567111

Cleves, P. A., Hart, J. C., Agoglia, R. M., Jimenez, M. T., Erickson, P. A., Gai, L., & Miller, C. T. (2018). An intronic enhancer of Bmp6 underlies evolved tooth gain in sticklebacks. PLOS Genetics, 14(6), e1007449. https://doi.org/10.1371/journal.pgen.1007449

Ellis, N. A., Donde, N. N., & Miller, C. T. (2016). Early development and replacement of the stickleback dentition. Journal of Morphology, 277(8), 1072–1083. https://doi.org/10.1002/jmor.20557

Ellis, N. A., Glazer, A. M., Donde, N. N., Cleves, P. A., Agoglia, R. M., & Miller, C. T. (2015). Distinct developmental and genetic mechanisms underlie convergently evolved tooth gain in sticklebacks. Development, (June), 2442–2451. https://doi.org/10.1242/dev.124248

Emera, D., Yin, J., Reilly, S. K., Gockley, J., & Noonan, J. P. (2016). Origin and evolution of developmental enhancers in the mammalian neocortex. Proceedings of the National Academy of Sciences of the United States of America, 113(19), E2617–E2626. https://doi.org/10.1073/pnas.1603718113

Erickson, P. A., Cleves, P. A., Ellis, N. A., Schwalbach, K. T., Hart, J. C., & Miller, C. T. (2015). A 190 base pair, TGFB responsive tooth and fin enhancer is required for stickleback Bmp6 expression. Developmental Biology, 401(2), 310–323. https://doi.org/10.1016/j.ydbio.2015.02.006

Erickson, P. A., Ellis, N. A., & Miller, C. T. (2016). Microinjection for Transgenesis and Genome Editing in Threespine Sticklebacks. Journal of Visualized Experiments, (111), e54055–e54055. https://doi.org/10.3791/54055

Farh, K. K. H., Marson, A., Zhu, J., Kleinewietfeld, M., Housley, W. J., Beik, S., … Bernstein, B. E. (2015). Genetic and epigenetic fine mapping of causal autoimmune disease variants. Nature, 518(7539), 337–343. https://doi.org/10.1038/nature13835

Fish, A., Chen, L., & Capra, J. A. (2017). Gene regulatory enhancers with evolutionarily conserved activity aremore pleiotropic than those with species-specific activity. Genome Biology and Evolution, 9(10), 2615–2625. https://doi.org/10.1093/gbe/evx194

Frazer, K. A., Pachter, L., Poliakov, A., Rubin, E. M., & Dubchak, I. (2004). VISTA: Computational tools for comparative genomics. Nucleic Acids Research, 32(WEB SERVER ISS.), W273–W279. https://doi.org/10.1093/nar/gkh458

Infante, C. R., Mihala, A. G., Park, S., Wang, J. S., Johnson, K. K., Lauderdale, J. D., & Menke, D. B. (2015). Shared Enhancer Activity in the Limbs and Phallus and Functional Divergence of a Limb-Genital cis-Regulatory Element in Snakes. Developmental Cell, 35(1), 107–119. https://doi.org/10.1016/j.devcel.2015.09.003

Jackman, W. R., & Stock, D. W. (2006). Transgenic analysis of Dlx regulation in fish tooth development reveals evolutionary retention of enhancer function despite organ loss. Proceedings of the National Academy of Sciences of the United States of America, 103(51), 19390–19395. https://doi.org/10.1073/pnas.0609575103

Jia, S., Zhou, J., Gao, Y., Baek, J.-A., Martin, J. F., Lan, Y., … Sucov, H. M. (2013). Roles of Bmp4 during tooth morphogenesis and sequential tooth formation. Development (Cambridge, England), 140(2), 423–432. https://doi.org/10.1242/dev.081927

Jones, F. C., Grabherr, M. G., Chan, Y. F., Russell, P., Mauceli, E., Johnson, J., … Kingsley, D. M. (2012). The genomic basis of adaptive evolution in threespine sticklebacks. Nature, 484(7392), 55–61. https://doi.org/10.1038/nature10944

Kandyba, E., Leung, Y., Chen, Y.-B., Widelitz, R., Chuong, C.-M., & Kobielak, K. (2013). Competitive balance of intrabulge BMP/Wnt signaling reveals a robust gene network ruling stem cell homeostasis and cyclic activation. Proceedings of the National Academy of Sciences, 110(4), 1351–1356. https://doi.org/10.1073/pnas.1121312110

Kawakami, K., & Shima, A. (1999). Identification of the Tol2 transposase of the medaka fish Oryzias latipes that catalyzes excision of a nonautonomous Tol2 element in zebrafish Danio rerio. Gene, 240(1), 239–244. https://doi.org/10.1016/S0378-1119(99)00444-8

Kobielak, K., Stokes, N., de la Cruz, J., Polak, L., & Fuchs, E. (2007). Loss of a quiescent niche but not follicle stem cells in the absence of bone morphogenetic protein signaling. Proceedings of the National Academy of Sciences, 104(24), 10063–10068. https://doi.org/10.1073/pnas.0703004104

Lee, C. S., Friedman, J. R., Fulmer, J. T., & Kaestner, K. H. (2005). The initiation of liver development is dependent on Foxa transcription factors. Nature, 435(7044), 944–947. https://doi.org/10.1038/nature03649

Levine, M., Cattoglio, C., & Tjian, R. (2014, March 27). Looping back to leap forward: Transcription enters a new era. Cell. Elsevier B.V. https://doi.org/10.1016/j.cell.2014.02.009

Lukoseviciute, M., Gavriouchkina, D., Williams, R. M., Hochgreb-Hagele, T., Senanayake, U., Chong-Morrison, V., … Sauka-Spengler, T. (2018). From Pioneer to Repressor: Bimodal foxd3 Activity Dynamically Remodels Neural Crest Regulatory Landscape In Vivo. Developmental Cell, 47(5), 608–628.e6. https://doi.org/10.1016/j.devcel.2018.11.009

Maurano, M. T., Humbert, R., Rynes, E., Thurman, R. E., Haugen, E., Wang, H., … Stamatoyannopoulos, J. A. (2012). Systematic localization of common disease-associated variation in regulatory DNA. Science, 337(6099), 1190–1195. https://doi.org/10.1126/science.1222794

O’brown, N. M., Summers, B. R., Jones, F. C., Brady, S. D., & Kingsley, D. M. (2015). A recurrent regulatory change underlying altered expression and Wnt response of the stickleback armor plates gene EDA. ELife, 2015(4). https://doi.org/10.7554/eLife.05290

Pispa, J., & Thesleff, I. (2003). Mechanisms of ectodermal organogenesis. Developmental Biology, 262(2), 195–205. https://doi.org/10.1016/S0012-1606(03)00325-7

Preger-Ben Noon, E., Sabarís, G., Ortiz, D. M., Sager, J., Liebowitz, A., Stern, D. L., & Frankel, N. (2018). Comprehensive Analysis of a cis-Regulatory Region Reveals Pleiotropy in Enhancer Function. Cell Reports, 22(11), 3021–3031. https://doi.org/10.1016/j.celrep.2018.02.073

Rebeiz, M., & Tsiantis, M. (2017, August 1). Enhancer evolution and the origins of morphological novelty. Current Opinion in Genetics and Development. Elsevier Ltd. https://doi.org/10.1016/j.gde.2017.04.006

Sabarís, G., Laiker, I., Preger-Ben Noon, E., & Frankel, N. (2019, June 1). Actors with Multiple Roles: Pleiotropic Enhancers and the Paradigm of Enhancer Modularity. Trends in Genetics. Elsevier Ltd. https://doi.org/10.1016/j.tig.2019.03.006

Sievers, F., Wilm, A., Dineen, D., Gibson, T. J., Karplus, K., Li, W., … Higgins, D. G. (2011). Fast, scalable generation of high-quality protein multiple sequence alignments using Clustal Omega. Molecular Systems Biology, 7(1), 539. https://doi.org/10.1038/msb.2011.75

Singh, D., & Yi, S. V. (2021). Enhancer Pleiotropy, Gene Expression, and the Architecture of Human Enhancer–Gene Interactions. Molecular Biology and Evolution, 38(9), 3898–3909. https://doi.org/10.1093/molbev/msab085

Square, T. A., Sundaram, S., Mackey, E. J., & Miller, C. T. (2021). Distinct tooth regeneration systems deploy a conserved battery of genes. EvoDevo, 12(1), 1–17. https://doi.org/10.1186/s13227-021-00172-3

Stepaniak, M. D., Square, T. A., & Miller, C. T. (2021). Evolved Bmp6 enhancer alleles drive spatial shifts in gene expression during tooth development in sticklebacks. Genetics. https://doi.org/10.1093/genetics/iyab151

Vainio, S., Karavanova, I., Jowett, A., & Thesleff, I. (1993). Identification of BMP-4 as a signal mediating secondary induction between epithelial and mesenchymal tissues during early tooth development. Cell, 75(1), 45–58. https://doi.org/10.1016/S0092-8674(05)80083-2

Wang, L., Siegenthaler, J. A., Dowell, R. D., & Yi, R. (2016). Foxc1 reinforces quiescence in self-renewing hair follicle stem cells. Science (New York, N.Y.), 351(6273), 613–617. https://doi.org/10.1126/science.aad5440

Wang, Y., Li, L., Zheng, Y., Yuan, G., Yang, G., He, F., & Chen, Y. (2012). BMP activity is required for tooth development from the Lamina to Bud Stage. Journal of Dental Research, 91(7), 690–695. https://doi.org/10.1177/0022034512448660

Xu, P., Balczerski, B., Ciozda, A., Louie, K., Oralova, V., Huysseune, A., & Gage Crump, J. (2018). Fox proteins are modular competency factors for facial cartilage and tooth specification. Development (Cambridge), 145(12). https://doi.org/10.1242/dev.165498

Xu, P., Yu, H. V., Tseng, K. C., Flath, M., Fabian, P., Segil, N., & Crump, J. G. (2021). Foxc1 establishes 1 enhancer accessibility for craniofacial cartilage differentiation. ELife, 10, 1–50. https://doi.org/10.7554/eLife.63595

